# Grp78 alleviates sodium iodate-induced retinal cell injury in vivo and in vitro

**DOI:** 10.1101/2021.03.22.436404

**Authors:** Jiang Shuang, Guo Yongpeng, Yi Ning, Li Hongdan, Liu Hua

**Author notes:** Correspondence to: Liu Hua; Li Hongdan. **E-mail:** Jiang Shuang.

## Abstract

**Objective:** Glucose-regulated protein 78 (Grp78) has been regarded as a main member of the endoplasmic reticulum proteins, Grp78 could protect cells from apoptosis under stress conditions. However, whether Grp78 could protect retinal pigment epithelium (RPE) cells from oxidative injury and then protect retinas from morphological changes and functional abnormalities remain undetermined. Here, we try to explore the effect of Grp78 on retinal cell injury induced by sodium iodate in vivo and in vitro.

**Methods:** To investigate whether Grp78 has a protective effect on RPE injury in vitro, human retinal pigment epithelium (ARPE-19) cells were treated with sodium iodate. The cell proliferation, morphology, apoptosis and ROS production assays were detected. In vivo, We established sodium iodate-induced retinal injury model in mice by intravenous injection of sodium iodate into tail vein. After that, we examined the morphology and function of retina in mice by fundus photography, OCT and ERG. Finally, we removed the retina of mice for histological examination.

**Results:** Grp78 significantly inhibited sodium iodate-induced reactive oxygen species (ROS), and decreased apoptosis of RPE in vitro. Furthermore, Grp78 significantly decreased the apoptosis of retinal cells in vivo, resulting in the inhibition of morphological changes of retina, and improving the function of retina. The underlying mechanisms included inhibited caspase3 and Nos, and increased expression of Bcl2, thereby protecting RPE from SI-induced ROS and apoptosis.

**Conclusion:** Grp78 could reduce the injury of retinal cells induced by sodium iodate in vitro and in vivo. These findings suggested Grp78 may become a new therapeutic target for retinal injury in clinical practice.

## Introduction

Age-related macular degeneration (AMD) is a degenerative disease of the retin leading cause of irreversible blindness in people over the age of 65 ^[1]^. In the industrialized world, AMD becomes one of the leading causes for visual impairment and blindness with no cure^[2, 3]^ . While the exact etiology of AMD is not known, numerous studies support there are many reasons for pathological changes including genetic factors, oxidative stress, inflammation and environmental actors^[4-6]^. Oxidative stress is considered the most important risk factor for AMD ^[7]^. Some studies have shown that endoplasmic reticulum stress plays an important role in the pathogenesis of AMD ^[8, 9]^ . For RPE cells are susceptible to oxidative stress induced by their own metabolic products or any other element, researchers try to lower the level of oxidative stress to reverse AMD.

Glucose-regulated protein 78(Grp78) is a member of the Endoplasmic Reticulum (ER) stress protein family and its major function is to fold and process the unfolded or malfolded proteins. Traditionally GRP78 regarded as an ER resident protein and act as a molecular chaperone in the lumen of ER^[10]^. Recently, several lines of evidence have demonstrated that GRP78 is a multifunctional protein and plays critical roles in the resistance to chemotherapy agents, proliferation, invasion and metastasis of many human cancers^[11]^. Under stress conditions, Grp78 could protect cells from apoptosis and maintain the cell viability. However, whether Grp78 protects retinal cells under stress is still unknown.

None of the models are true AMD, yet their utility lies in the fact that they recapitulate specific aspects of the disease. The sodium iodate model is widely used to study the molecular mechanism of cell injury in AMD as it represents the diseaseassociated increase in oxidative stress and induces consistent and selective damage to the RPE^[12, 13]^. Exposure to sodium iodate results in a primary death of the RPE followed by a secondary death of the overlying photoreceptors, similar to what is observed in advanced atrophic AMD^[14-17]^. A major outstanding problem in AMD is that sodium iodate can induce consistent and selective injury to the RPE^[18]^.

In this study, we use sodium iodate to induce retinal cell injury in vitro and in vivo. In the article, we identify whether Grp78 could reduce the retinal cell injury induced by sodium iodate in vitro and in vivo. Grp78 could reduce the injury of retinal cells induced by sodium iodate in vitro and in vivo. These findings suggested Grp78 may become a new therapeutic target for retinal injury in clinical practice.

## Materials and Methods

### Cell line and reagent

Human RPE (ARPE-19) cells were purchased from American Type Culture Collection (ATCC, Manassas, VA, USA) and maintained in DMEM/F12 medium containing 10% fetal bovine serum (FBS; Hyclone, Shanghai, China) and 1% penicillin/streptpmycin.

### Cell viability assay

In the first experiment, we treated ARPE-19 cells with different concentrations of SI (0μM, 100μM, 300μM, 500μM, 800μM, 1000μM) for 24 hours and 48 hours. ARPE-19 cells were cultured at 5,000 cells per well in 96 -well tissue culture plates. After 24 h after plating, cells were washed 3 times with PBS and then treated with the sodium iodate for 24h and 48h in DMEM/F12 medium containing 1% FBS. At the end of the culture period, cells were washed with ice cold PBS, the MTT reagents (5mg MTT in 10ml PBS) were added according to the manufacturer’s instructions and the absorbance was measured at 490 nm using a microplate reader. Mean values were calculated from three independent experiments. In the second experiment, we treated ARPE-19 cells with SI(500μM) for 48 hours. In the third experiment, ARPE-19 cells were pretreated with Grp78 protein (0, 0.1, 1, 2μg/ml) for 2 hours, and then were treated with SI(500μM). Cell viability was measured using the above described methods .

### Cell apoptosis assay

Hochest 33342 staining was performed after pre-treatment of ARPE-19, and apoptotic cells were identified based on the morphological changes in the nuclear assembly involving chromatin condensation and fragmentation. Flow cytometry was also used to detect apoptotic cells. Cell apoptosis was measured by Annexin-V FACS according to the manufacturer’s protocol (Calbiochem). Briefly, after treatment, cells were washed twice with PBS and incubated in 300μl binding buffer containing 3μl annexinV–FITC and 3μl of propidium iodide in the dark for 15 min at room temperature. The stained samples (containing 200,000 cells/sample) were then analyzed on a FACS Calibur flow cytometer within 1h following the manufacturer’s protocol (BD Biosciences).

### Measurement of ROS production

ARPE-19 cells were pretreated with Grp78 protein (0, 0.1, 1, 2μg/ml) for 2 hours, and then were treated with SI(500μM). ROS production was determined by carboxy-H2DCF-DA (Calbiochem) staining assay. ARPE -19 cells were incubated with 1μM carboxy-H2DCF-DA at 37°C for 30min. Cells (1×10^6^) were then re-suspended in PBS and analyzed by flow cytometry. The percentage of fluorescence-positive cells was recorded on a FACS Calibur flow cytometer (BD Biosciences) using excitation and emission filters of 488 and 530nm.

### Animals

Six to eight weeks old male C57BL/6J mice (Beijing Weitong Lihua animal Co. ltd) were used in the experiment. The mice were housed in a standard laboratory environment and maintained on a 12-hour light–dark cycle at 21°C. This study was carried out in strict accordance with the recommendations in the Guide for the Care and Use of Laboratory Animals of the National Institutes of Health. The protocol was approved by the Committee on the Ethics of Animal Experiments of the Jinzhou Medical University (Protocol Number: JYDSY-KXYJ-IEC-2019-002). All surgery was performed under pentobarbital sodium anesthesia, and all efforts were made to minimize suffering.

### Animal experimental setting

In the experiment of defining the concentration of sodium iodate, sixty mice were randomly divided into six groups: control (n=10), SI (10mg/Kg, n =10), SI (20mg/Kg, n = 10), SI (30mg/Kg, n=10), SI (45mg/Kg, n=10) and SI (60mg/Kg, n =10). Sterile 1% sodium iodate solution was freshly prepared from solid sodium iodate (Solarbio) diluted in 0.9% sterile phosphate-buffered saline (PBS). Mice were anesthetized with 1% pentobarbital sodium (5ml/kg body weight). Mice from the experimental group were injected via the tail vein with sodium iodate (10, 20, 30, 45, 60mg/kg). Control mice were injected with similar volumes of PBS instead of sodium iodate. Mice were sacrificed at 1 -7 days after injection. In the experiment of mice pretreated with Grp78, thirty mice were randomly divided into two groups: SI+PBS (n=15), SI+Grp78 (n = 15). Before intravenous injection of sodium iodate (30mg/Kg), Grp78(1µg/µl) or PBS were injected into the eyeballs of mice.

### Grp78 intravitreal injection

Before tail vein injection, Tropicamide phenylephrine eye drops (Santen) and Oxybuprocaine Hydrochloride eye drops (Santen) were administered until the pupils of the mice dilated. With a surgical microscope, the needle of a microsyringe was inserted into the vitreous cavity 0.5 mm posterior to the corneoscleral limbus meanwhile avoiding the crystalline lens . 2 µl of Grp78 (1µg/µl) was injected into SI+Grp78 group and the same dose of PBS was injected into SI+PBS group. Following injection, ofloxacin eye ointment (Shenyang Xingqi Pharmaceutical Co., Ltd., Shenyang, China) were administered for preventing infection.

### SD-OCT and Fundus Photography

Ultrahigh-resolution SD-OCT and Fundus images were obtained on live anesthetized animals using the Micron I V retinal imaging system (Phoenix Research Laboratories, Pleasanton, CA). Conventional mydriasis and corneal protection are performed in accordance with the above method. The positions of retina in SD-OCT were measured 220μm outward from the center of optic disk and the optic disk level. Optical coherence tomography was performed on days 0,1,3,5 and 7 to monitor the retinal morphology.

### Electroretinography (ERG)

Mice were dark-adapted overnight before analysis. Pupils were dilated with tropicamide phenylephrine eye drops (Santen). Mice were anesthetized with 1% pentobarbital sodium and placed on a temperature-controlled workingplatform at 37°C. Contact electrodes were placed on the corneal surface and visual responses were recorded with ERG system (Ai Erxi, Chongqing, China). Flash responses were obtained. The a-wave amplitude was measured from the baseline to the trough of the a-wave, while the b-wave amplitude was measured from the trough of the a-wave to the peak of the b-wave.

### Fluorescein fundus angiography

General fundus photography and fluorescein angiography were performed with a commercial camera and imaging system (Micron IV, Phoenix research labs, Pleasanton, CA). The mice were intraperitoneally anesthetized by 1% pentobarbital sodium in mice (6ml/kg) and intraperitoneally injected with 40 µl of 2% fluorescein sodium (Akorn, NDC17478-253-10) per 10 g body weight.

### Western blot analysis

For extraction of total cellular protein, cells were lysed in RIPA buffer with PMSF. Protein concentration was quantified using the BCA kit (Pierce Biotechnology, Inc., Rockford, IL). Proteins were separated and transferred to PVDF membranes. The membranes were incubated overnight at 4°C with the caspase-3, Bcl2, Bad, Nos and β-actin (1:1000) (Cell signaling technology, Danvas, MA). Thereafter, the membranes were incubated with HRP-labeled anti-rabbit secondary antibodies (1:1000) for 1h at room temperature. At last, the membrane was visualized by enhanced chemiluminescence kit (Thermo Fisher Scientific Inc., Rockford, IL, USA). The intensity of the bands was measured with a scanning densitometer (Bio-Rad, Shanghai, China) coupled with Bio-Rad analysis software.

### Statistical analysis

Each experiment was repeated a minimum of three times, the mean value of the repetitions was calculated, and this value was used in the statistical analysis. Results were presented as mean ±SD. Values of ERG and OCT were analyzed by 2-tailed Student *t*-test and one-way ANOVA . Nonparametric Kruskal-Wallis (K-W) test was used for western blotting analysis. A P-value of <0.05 was considered statistically significant.

## Results

### 1. Sodium iodate induced ARPE-19 cells injury in vitro

To explore which concentration of sodium iodate is suitable for RPE cells injury, MTT assay was chosen to detect the IC50 (the half maximal inhibitory concentration is a measure of the effectiveness of sodium iodate in inhibiting cell viability) of sodium iodate-induced RPE cells injury. ARPE-19 cells were stimulated with increasing concentrations of sodium iodate (0, 100, 300, 500, 800, 1000μM) for 24h and 48h, and their viabilities were measured by MTT assay. The results shown in Fig.1A demonstrated that sodium iodate depressed ARPE-19 cells viability in a dose-dependent manner. The IC50 of sodium iodate is about 500μM at 24h and 312.67μM at 48h; (Fig.1A). Fig.1B showed sodium iodate reduce cell viability in a time-dependent manner. With time extending, the viability of ARPE-19 cells came down in 500μM sodium iodate. Fig.1C and 1D showed the nucleus and cell morphology of ARPE-19 treated with different concentrations of sodium iodate. Hochest 33342 staining showed that apoptotic nucleus increased significantly in a time-dose-dependent manner in Fig.1C. So, 500μM was chosen for the following experiments . Western blot showed that in 500μM sodium iodate, the Grp78, Nos, and Caspase 3 expressions increased in a time dependent manner (Fig. 1E).

**Fig 1.**
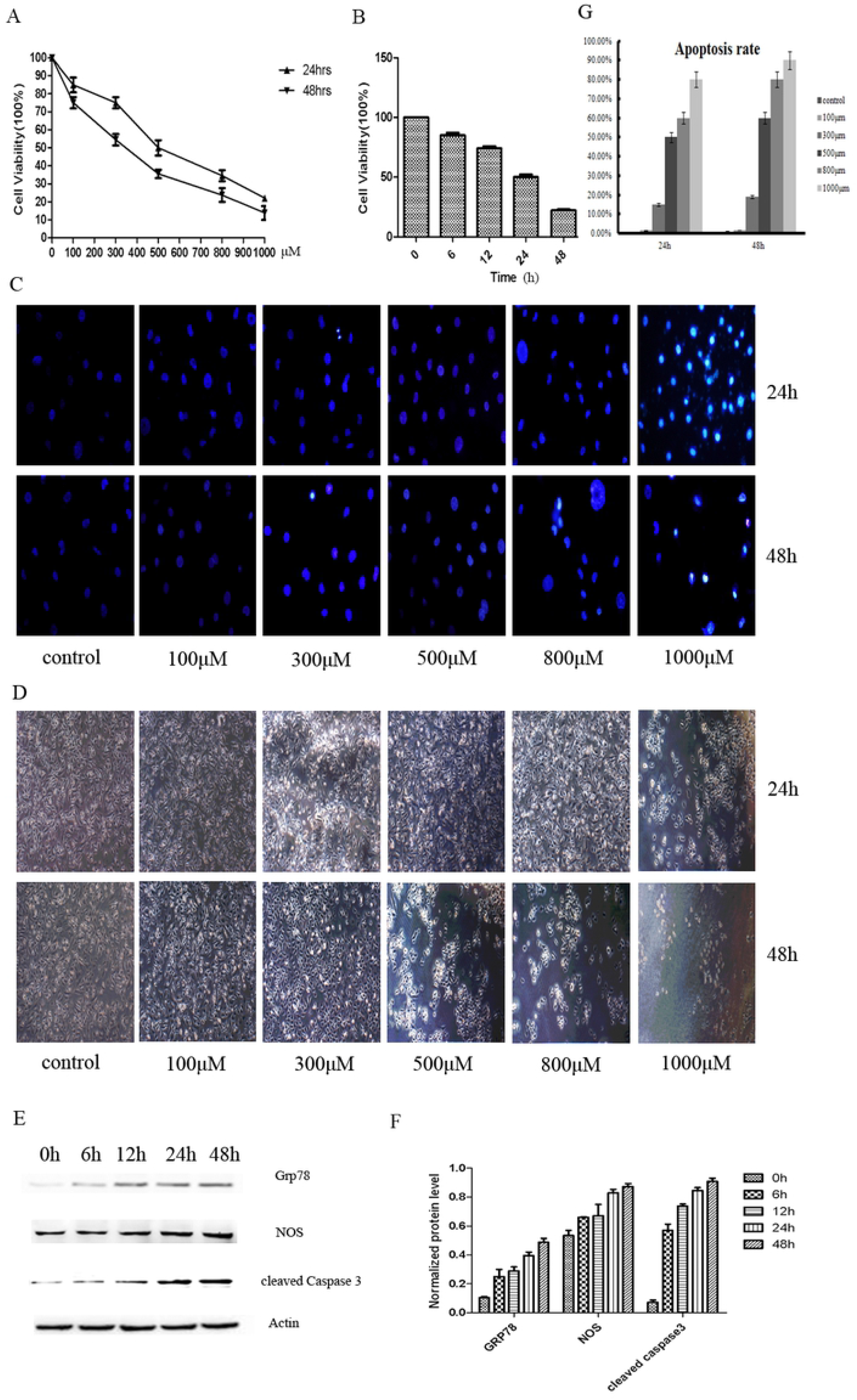
Sodium iodate induced ARPE-19 cell injury in vitro. A demonstrated that sodium iodate depressed ARPE-19 cells viability in a dose-dependent manner. The IC50 of sodium iodate is about 500μM at 24h and 312.67μM at 48h. B showed that showed that the decrease of ARPE-19 cell viability induced by sodium iodate at 500μM concentration was time-dependent. C showed the apoptotic nucleus of ARPE-19 cells by Hochest 33342 assay. D showed the morphology of RPE cells treated with sodium iodate after 24h and 48h. E showed SI could promote the expression of Grp78, Nos, Caspase 3 in 48h by WB.

### 2. Sodium iodate induced retinal injury in vivo

To ensure which concentration of sodium iodate is suitable for retinal injury model, Optical coherence tomography (OCT) and color fundus photographs of mice treated with different doses of SI were used to detect the changes of retinal. Tab.1 showed the proportion of retinal damage after different concentrations of SI injection. From Fig.2, we can see that 30mg/kg body weight is the lowest concentration to retinal injury model. OCT and color fundus photographs showed that no obvious changes were found for the low dose(10 mg/kg and 20 mg/kg) group at any time point compared with the controls (Fig. 2). Middle dose (30 mg/kg) was the minimum concentration that can cause retinal damage. Higher concentrations(45 and 60mg/kg) can lead to rapid changes in the retina. Retinal vasculature and optic nerve tissue appeared to remain normal at all time points and concentrations. The histologic changes in paraffin sections showed that starting from day 3, the RPE injury appeared (Fig.2B). Histological changes in paraffin sections of the retina showed that RPE damage began on the third day, and significant loss of RPE cells and thinning of outer nuclear layer of retina (ONL) were detected on the fifth and seventh days. The histologic changes were in accordance with OCT and color fundus photographs. To sum up, we selected 30mg/kg body weight as the optimal concentration at the followed experiments.

**Fig 2.**
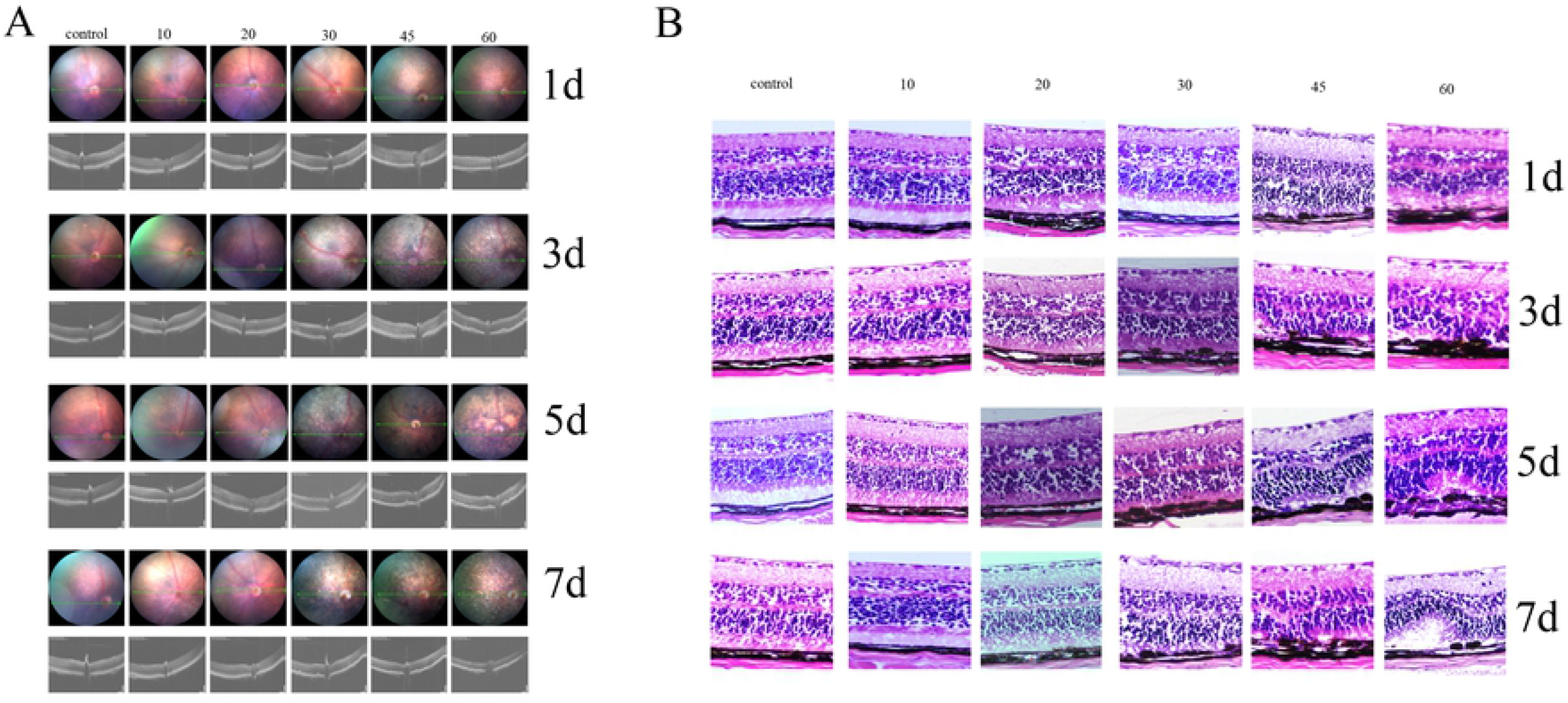
Sodium iodate induced retinal injury in vivo. A:The OCT and color fundus photographs showed that no obvious changes were found for the low doses (10 and 20 mg/kg) group compared with the control group at the same time period . Middle dose (30 mg/kg) is the lowest SI concentration for retinal damage . Higher doses (45 and 60mg/kg) were associated with progressive retinal degeneration characterized by significant pigmentary changes of the RPE layer at 3 day. Retinal injury in mice occured on the first day, when the concentration of SI more than 30 mg/kg. And the fundus of the mice were covered with yellow and white disease similar to “drusen”. B:The HE staining of the retina after injection of sodium iodate. The histologic changes in paraffin sections of the retina after injection of sodium iodate (30, 45 and 6 0mg/kg) showed the disordered arrangement of RPE cells at 1 day and the thinning of photoreceptor cell layers at 5 day.

### 3. Grp78 inhibited sodium iodate-induced ARPE-19 cell injury in vitro

The result in Fig.3A and 3B showed that Grp78 pretreatment (0μg/ml, 0.1μg/ml, 1μg/ml and 2μg/ml) significantly inhibited apoptosis rate of ARPE-19 cells treated with sodium iodate (500μM). Flow cytometry analysis also confirmed the protective effect of GRP78 on RPE cells(Fig.3C). The apoptosis rate of ARPE-19 cells treated with 500μM sodium iodate were significantly increased compared to the control. However, Grp78 can reduce the apoptosis rate significantly in a dose-dependent manner (Fig.3D). Compared with the group treated with sodium iodate, GRP78 treatment group (1 ug/ml and 2 ug/ml) had significant statistical differences (*P*<0.01, Fig.3E and 3F). To detect the oxidative stress level, we examed the expression of Reactive Oxygen Species (ROS) by ROS kit and Nitric Oxide Synthase (NOS) by Western Blot. Treatment with 500μM sodium iodate induced a significant increase in ROS to more than one fold for the control and Grp78 could inhibit the oxidative stress level (*P*<0.05, Fig.3G, Fig.3H and Fig.3I). To confirm the possible mechanism of Grp78 on ARPE-19 cell apoptosis, western blot analysis was performed to detect the expression levels of Caspase 3, Bcl-2 and Bad in ARPE-19 cells induced by sodium iodate at concentration of 500μM. The western blot results in Fig. 3H showed that Grp78 inhibited the expressions of caspase-3 and Bad, and promoted the expression of Bcl-2. All the above indicated that the protective effect of Grp78 might be due to anti-apoptosis of ARPE-19 after SI treatment.

**Fig 3.**
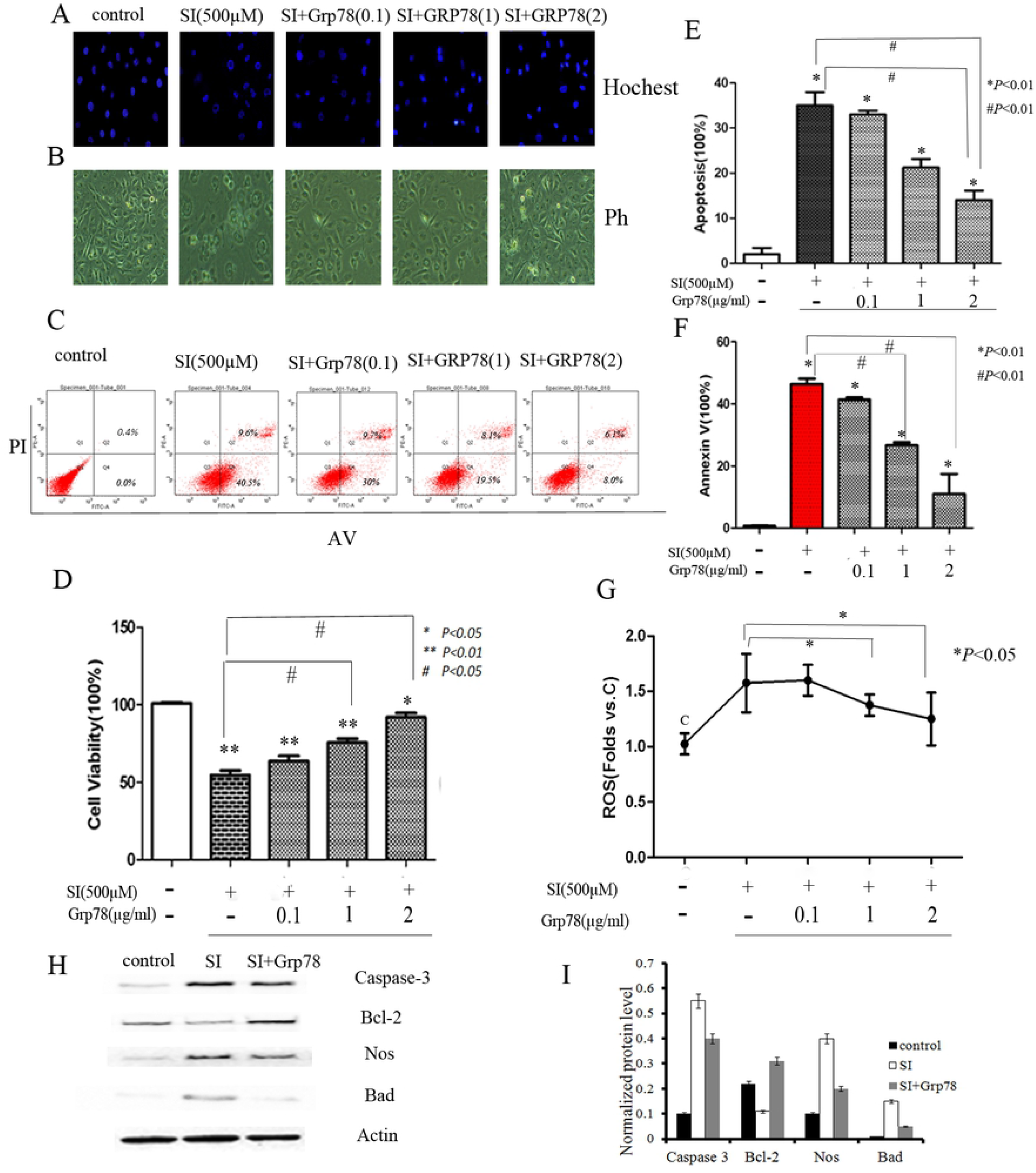
Grp78 inhibited sodium iodate induced ARPE-19 cell injury in vitro. A and E: The result showed that Grp78 pretreatment (0μg/ml, 0.1μg/ml, 1μg/ml and 2μg/ml) significantly inhibited the number of ARPE-19 cell apoptosis rate treated wi th sodium iodate (500μM). B: Cell morphology showed Grp78 pretreatment (0μg/ml, 0.1μg/ml, 1μg/ml and 2μg/ml) significantly inhibited ARPE-19 cell apoptosis. C and F: Grp78 inhibited the ARPE-19 cell apoptosis by flow cytometry assay. Fig.3F is the statistical analysis of apoptotic rate. D: MTT showed the cell viability of ARPE-19 cell treated with 500μM SI were significantly decreased compared to the control. And Grp78 could increase the cell viability significantly (* ^/#^ *P*<0.05,** *P*<0.01). G: ROS assay showed the protective effect of Grp78 in ARPE-19 cell injury caused by sodium iodate. (* *P*<0.05) H and I: WB showed protein levels of Caspase 3, Bcl-2, Nos and Bad in the ARPE-19 cell treated with SI and SI+Grp78. The result showed that Grp78 inhibited the expressions of caspase-3,Nos and Bad, and promoted the expression of Bcl-2.

From the figures above, we can see that with the increase of treatment concentration of Grp78, the protective effect is significant enhanced. Together, Grp78 inhibits sodium iodate-induced ARPE-19 cell apoptosis and oxidative stress in vitro.

### 4. Grp78 inhibits sodium iodate-induced retinal injury in vivo

Grp78 significantly inhibits sodium iodate-induced retinal injury. After the intravitreal injection of Grp78 (0.1, 1 and 2µg/ml), compared with the sodium iodate group, the fundus changes were lighter and slower. The lesions appeared later and lighter than sodium iodate group. On the first day post-sodium iodate injection, there was no obvious changes in fundus photograph and OCT (Fig.4A). On the third day post-sodium iodate injection, the fundus photographs images appeared to some pigment changes while there was not obvious pathological change in OCT. Subsequently, the fundus lesions became more, but were less than the contemporaneous sodium iodate group. After sodium iodate injection 7 day, the retinal structure could be identified in the Grp78 treatment group, but the retinal structure in sodium iodate group was still disordered (Fig.4A). The fundus photographs images on the first day post-sodium iodate injection appeared to obvious pigment change,while there were no observable changes in OCT image. On the third day post-sodium iodate injection, the outer nuclear layer (ONL) which is hypo-reflective in the normal retina became hyper-reflective and completely indistinguishable from the outer plexiform layer (OPL). Meanwhile, the retina of mice demonstrated loss of normal inner/outer segments (IS/OS) architecture, though the external limiting membrane (ELM) appeared intact. In the fundus photography of the same period, the fundus appeared gray white and the RPE pigment changes increased. On the fifth day post-sodium iodate injection, the hyper-reflective zones extended between the ONL and the RPE layer are mostly in OCT image, and the optic disc became pale in fundus photographs images. The change of the 7th day was more obvious than before: prominent loss of the RPE, ONL and IS/OS layers. The OCT image taken on the eighth day showed irregularly shaped patterns of hyper-reflectivity with granular texture at the site of the RPE. Furthermore, the ONL was markedly thin and irregular in shape at this time point. The lesions of RPE pigment change became smaller than before, but the scope of the lesion was almost spread throughout the entire retina (Fig.4A).

Grp78 significantly inhibits sodium iodate-induced retinal cell apoptosis and lipid deposition in RPE cell. To further investigate the structural changes in OCT and fundus photographs, we examined the histologic changes in paraffin sections (Fig.4B). On the fifth day post-sodium iodate injection, massive disruption of the RPE, IS/OS and focal reduction of photoreceptor cells produced irregular profiles of folding in the outer retinal layer. On the 7th day post-sodium iodate injection, the retinal thickness became further thinner. The structure of the RPE layer was not clear, and almost no RPE cells were noted (Figure 4B). The retinas of Grp78 group were slightly thinner than those of PBS group, but thicker than retinas in sodium iodate group. The cellular stratification of the retinas in Grp78 group was neater (Figure 4B). Quantification of nucleus of the ONL and IS/OS confirmed the protection of Grp78 on retina injury induced by sodium iodate.

**Fig 4.**
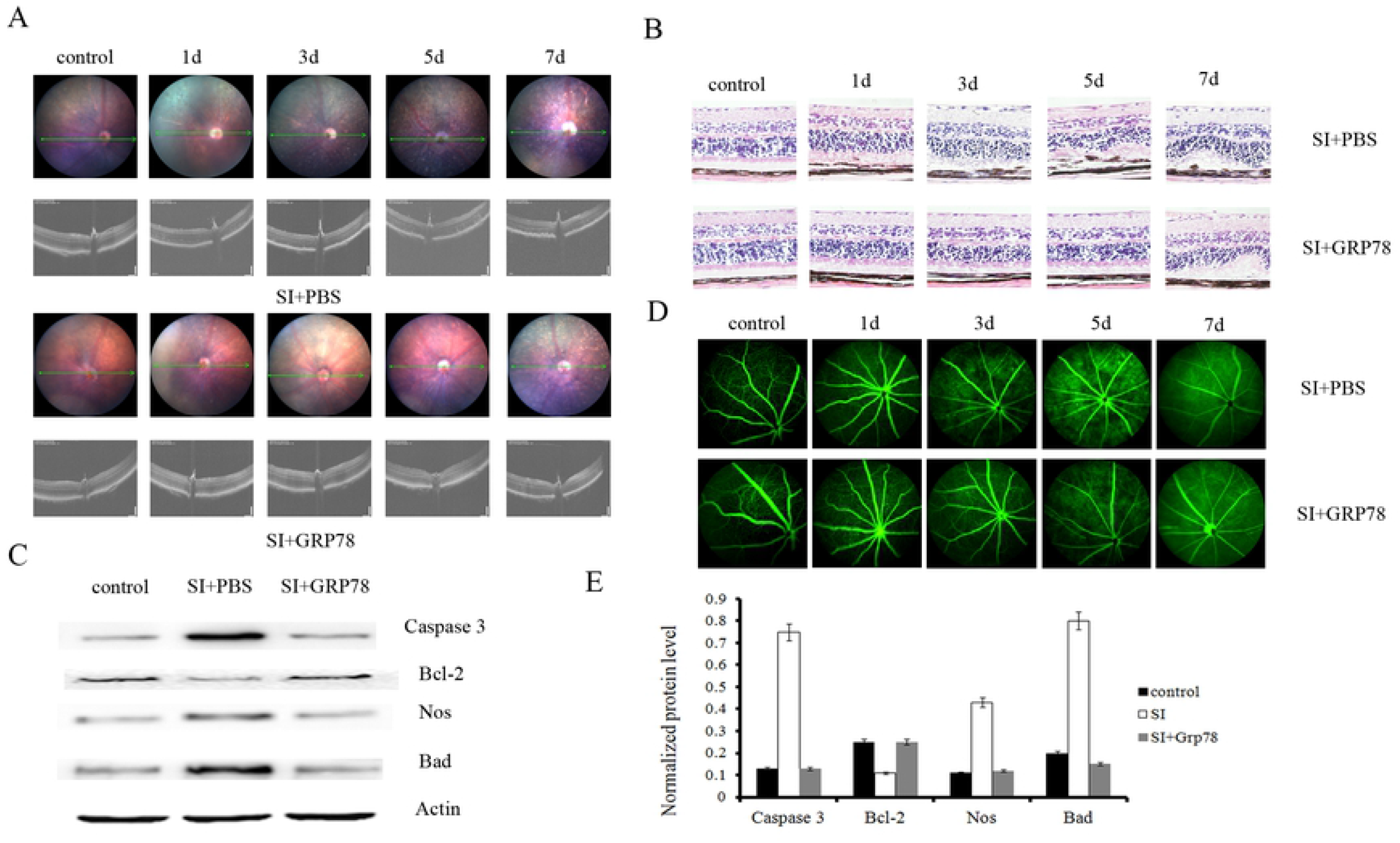
Grp78 inhibited sodium iodate induced retinal injury in vivo. A: The OCT and color fundus photographs showed that the injury of SI+GRP78 group is latter and lighter than the SI+PBS group before 7 day. B: HE staining showed the injury of RPE cell layer of SI+ GRP78 group is latter and lighter than the group which treated with SI+PBS before 7 day. C and E: Western blot of retina protein lysates showed protein levels of Caspase 3, Bcl-2, Nos and Bad in the control, SI and SI+ GRP78 groups. D: Fluorescein fundus angiography showed that the vascular leakage of SI+Grp78 was less than the vascular leakage of SI+PBS before day 7.

To better characterize the results, we used WB assay to detect the expressions of Caspase 3, Bcl-2, Nos and Bad. The results showed that Grp78 could inhibit the apoptosis after injection of sodium iodate in vivo (Figure 4C and E).

To explore the vascular leakage, fluorescein angiography was selected to show the difference in groups (Figure 4D). The results showed that Grp78 could inhibit the vascular leakage compare with the PBS group in some extent.

### 5. Grp78 improved sodium iodate-induced retina function degeneration

Retinal function changes were examined by flash ERG at 1, 3, 5, and 7 days post-injection. Analysis of both a- and b-waves showed that Grp78 pre-treatment dramatically up-regulated the amplitude of the waves on the 1, 3, 5, 7 days (Figure 5) and in a time-dependent manner. The intravitreal injection of Grp78 partially improved the retinal function, implying that it could protect or rescue the visual function of sodium iodate-treated mice.

**Fig 5.**
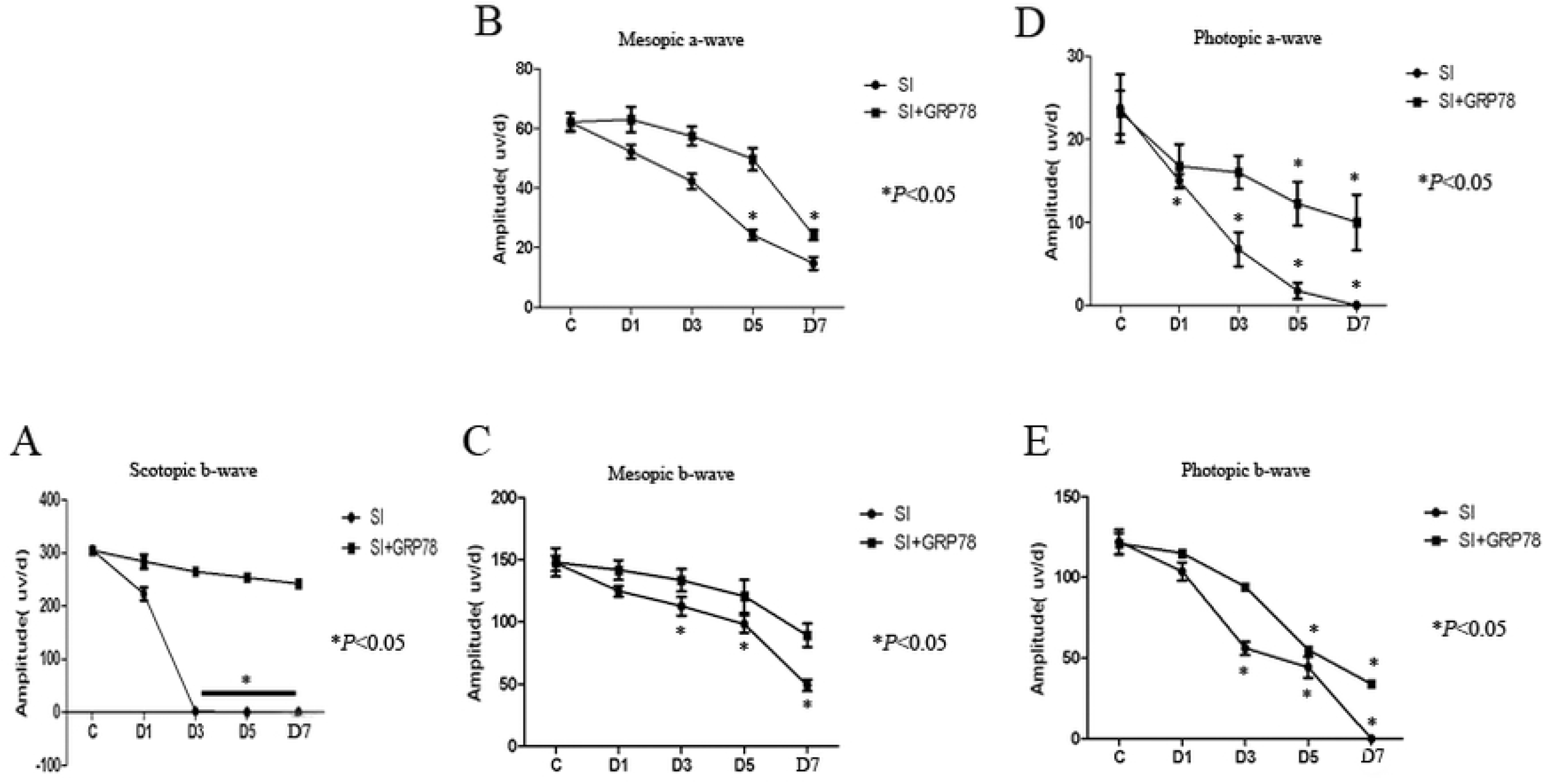
Grp78 improved sodium iodate induced retina function degeneration. A: The b-wave amplitudes of SI+GRP78 injection in scotopic (0.003cd.s.m ^-2^) was progressively, but at 3 day, the group of SI+PBS injection was elicited no response anymore. However, in the SI+GRP78 group, the b-wave amplitudes still remained 241.75±15uv/d at 7 day. It showed that GRP78 can alleviate the retinal injury of SI induced effectively. B, C a- and b-wave amplitudes in mesopic (3cd.s.m^-2^) was progressively, but the a-and b-wave amplitudes of SI+PBS group still lower than the SI+GRP78 group. It showed that GRP78 can alleviate the retinal injury induced by SI effectively. D,E a- and b-wave amplitudes in photopic (10cd.s.m^-2^) was progressively, a-and b-wave amplitudes of SI+PBS group still lower than the SI+GRP78 group. On the seventh day, almost no amplitude could be detected in SI+PBS group. However, the amplitude of SI+GRP78 group still remain 10(±3)uv/d and 34(±6)uv/d. It showed that GRP78 can alleviate the retinal injury induced by SI effectively.

## Discussion

In this study, we first explored the concentration of sodium iodate, which causes retinal cell damage in vivo and in vitro. We found that Grp78 can inhibite SI-induced ROS in RPE, and decreased RPE cell apoptosis in vitro and retinal cell apoptosis in vivo. To the best of our knowledge, this is the first study that reported Grp78 in the prevention of SI-induced RPE and retinal injury.

By western blot assay, we found that Grp78 could reduce the expression of caspase 3 and increase the expression of Bcl 2. Further, we detected the ROS level of ARPE-19 cells treated with sodium iodate. The level of ROS production is significantly increased after treatment of sodium iodate, indicating oxidative stress as a result of sodium iodate . After pretreatment with Grp78, we observed that Grp78 can retard this trend in a certain extent. Some scholars had also used different drugs to control the increase in ROS products to alleviate AMD . Our results demonstrate that sodium iodate activates caspase 3-mediated cell death in the RPE. Some scholars believed that cell death caused by SI is mediated by FasL^[19]^. Grp78 significantly reduces the extent of this cell death induced by SI. These findings suggest that Grp78 can protect RPE cells from oxidative stress in vitro.

Based on the protective effect of Grp78 to ARPE-19 in vitro, we tried to confirm whether Grp78 have the same effect in vivo. After injecting sodium iodate into rat tail vein ,the changes of “drusen”, pigment change, atrophy or even disappearance of RPE cells, and photoreceptor cells injury in retinas of mice were detected through the fundus photography or OCT. Further, to verify the role of Grp78 in the process, Grp78 pure protein was injected intravitreously before sodium iodate injection. The impact on retinas of Grp78 showed significant protective effect both in morphology, function and paraffin section staining. Although we achieved significant protection by Grp78, the effet was not total. One potential explanation is that the delivey of Grp78 was not ideal. This may not have resulted in optimal drug levels in the retina for protection against SI-induced oxidative stress. Future work would be to analyze and optimize the concentration of Grp78 and timing of delivery required to achieve maximal protection.

Results showed that sodium iodate can lead to tremendous retinal morphological changes, mainly for similar drusen deposition which showed as high reflection area in OCT. In the eighth days after injection, RPE layer almost disappeared, which was consistent with the results of H E staining. After administration of Grp78, “drusen” was significantly less than that of the same period in sodium iodate group. These results suggested that Grp78 can effectively improve the injury of retinal morphology induced by sodium iodate. In view of our short observation time, we will make a longer observation in the next experiment.

After the injury of the retina, the functional changes appeared earlier than the morphology. ERG is an objective and quantitative detection method, which is the most sensitive indicator of the response function of the retina, and is widely accepted by many scholars ^[7, 20]^. The results showed that the amplitudes of both a- and b- waves decreased, indicating that sodium iodate caused injury to the inner and outer layers of the retina. We observed the effect of Grp78 on the retina by recording the amplitude of a- and b-wave. The results showed that the amplitude of a-, b-wave in different stimulus intensities in Grp78 group were higher than those of sodium iodate group at different time points, which illustrated that Grp78 has certain protective effect on the function of retina. This study shows that Grp78 can effectively improve the function of photoreceptors, Müller cells and bipolar cells in the retina.

In summary, our study confirmed Grp78 preserves SI-induced RPE and retinal injury by inhibiting ROS and decreasing apoptosis in vivo and in intro. The underlying mechanisms include the activation of Bcl2 and the inhibition of caspase3. As such, we suggest that Grp78 is a potential target for the prevention and treatment of AMD. This study also provides strong support for the view that intravitreal injection therapy may play an important role in future therapeutic strategies.

## Grant Supports

This article is financially supported by Science and Technology Project of Liaoning province (2019-BS-101), Doctoral Foundation Project of Liaoning Science and Technology Department and Ao Hong Bo Ze Pharmaceutical Innovation Foundation, the president Fund of Jinzhou Medical University(XZJJ20130213) .

## Conflict of interest

The authors have declared that no competing interests exist.

**Table 1: Percentage of mice successfully injected with sodium iodate having retinal injury.**

